# Genomes of *Betacoronavirus gravedinis* from white-footed mice in New York City and a phylogenetically weighted model of its probable distribution in North America

**DOI:** 10.64898/2026.06.30.735598

**Authors:** Benjamin Kaza, Michael D. Catchen, Giulia B. de Gennaro, Jordan D. Zehr, Marie V. Lilly, Laura D. Plimpton, Maria Diuk-Wasser, Chloe M. Murrell, Alivia Ishee, Laura B. Goodman, Gary R. Whittaker, Amandine Gamble, Ximena Olarte-Castillo

## Abstract

Rodents are an important reservoir of zoonotic viruses and are ubiquitously present in densely populated urban areas. *Betacoronaviruses* in the *Embecovirus* lineage are well known to infect both humans and animals and have established rodent reservoirs. Here three *Betacoronavirus gravedinis* genomes were sequenced and characterized in white footed mice *(Peromyscus leucopus*, commonly white footed mice) collected in New York City, the second most populous city in North America. The genomes were distinct from mouse hepatitis virus (MHV), the prototype mouse betacoronavirus, and highly similar and identical in one case to previously characterized *B. gravedinis* sequences from white footed mice in Connecticut. Codon aware evolutionary models were used to identify specific sites under positive selection within the spike protein of *B. gravedinis*. A novel method was developed to predict the probable geographic distribution of the virus using publicly available data from the Global Biodiversity Information Facility to generate a weighted distribution map highlighting overlapping potential host ranges based on the evolutionary distance using a high resolution cytocrome B (CYTB) phylogeny of rodent species with potentially overlapping ranges. Our models predict three current hotspots of circulation in North America under different possible transmission regimes, and an additional fourth hotspot was predicted to arise in a warming future. This study highlights the continued need for biodiversity-informed surveillance of potential zoonotic pathogens in rodents.

## INTRODUCTION

Emerging infectious diseases of zoonotic origin continue to threaten public health globally, with most emerging human pathogens originating in animals^2^. The COVID-19 pandemic caused by the zoonotic betacoronavirus SARS-CoV-2, underscores the importance of continued bio surveillance of wildlife that are likely to harbor viruses that are capable of causing outbreaks in humans^4–7^. While betacoronaviruses are often associated with bat reservoirs, the *Embecovirus* subgenus is often associated with rodent hosts. *Embecoviruses* include mouse hepatitis virus (*Betacoronavirus muris*), as well as two circulating seasonal human viruses HCoV-OC43 (*Betacoronavirus gravadinis*) and HCoV-HKU-1 (*Betacoronavirus hongkongense*). The *B. gravadinis* species includes bovine coronavirus (BCoV), equine coronavirus (ECoV), canine respiratory coronavirus (CRCoV), and porcine hemagglutining encephalitis virus (PHEV), as well as a human enteric coronavirus strain 4408^8^. As such *B. gravadinis* includes a wide diversity of animal, human and zoonotic viruses and has been previously described as an example of a “promiscuous” coronavirus, seemingly capable of infecting many mammalian species^9^.

The sequencing and phylogenetic characterization is reported herein of three additional *B. gravedinis* genomes in *Peromyscus leucopus* (white footed mice), a widespread North American rodent species known for its ecological overlap with humans and its role as a reservoir for important zoonotic pathogens such as hantavirus and *Borellia burgdorferi* (the Lyme disease spirochete)^10–12^. Using metagenomic sequencing and genome-resolved phylogenetics, these genomes are proposed to be members of *B. gravedinis* in the *Embecovirus* subgenus clustering with the recently discovered *Peromyscus* coronavirus (PCoV) genomes identified in Connecticut^13^. As with all previously reported members of the *Embecovirus* subgenus, the genomes contains a hemagglutinin-esterase (HE) gene that are thought to bind O-acetylated sialic acid residues^14^. The spike protein resembles that of porcine hemagglutinating encephalitis virus (PHEV), which contains both a functional furin cleavage site (RRSRR|S) and a fusion-activating S2’ (R|S) site adjacent to the conserved fusion peptide^15^.

In the absence of virus isolation, we developed weighted Species Distribution Models using the phylogenetic distance from *P. leucopus* as a surrogate metric of the likelihood of re-sequencing the same or similar genomes in other North American rodent species^16^. In order to account for the oftentimes unpredictable relationship between phylogenetic distance and ability to host a particular virus^17^, we generated models using multiple hypotheses of how this virus may be transmitted to predict where these genomes are likely to be found on the continent building off of existing modeling frameworks from plants for inferring host infectability from phylogenetic signal^18,19^

## RESULTS

### Phylogenetic analysis and taxonomy

Whole-genome sequences were obtained of the two 2023 samples (SM009 and SM004). For the 2024 sample (SM074), NSP12 and spike (S) gene sequences were obtained. A whole-genome comparison of the two sequences obtained in this study (30,882 nucleotides, nt) revealed they are 99.9% similar and 97.7% similar to three Betacoronavirus genomes previously reported in white-footed mice in Connecticut. Phylogenetic tree of the conserved NSP12 gene and the spike gene showed that the three *B. gravedinis* genomes detected in this study clustered with these viruses (Figure 1).

**Figure 1:**
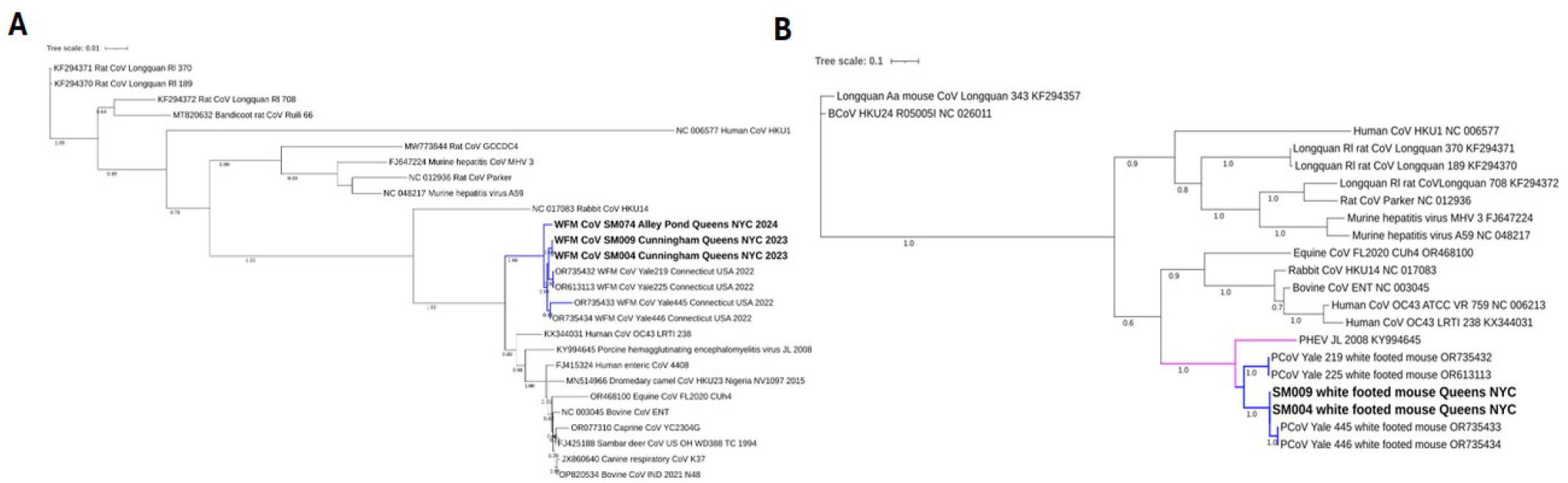
Phylogeny of the RdRp gene (a) and Spike (b)

The viruses are provisionally within the species *B. gravedinis* based on phylogenetic placement and putative genome organization consistent with established members of this clade. In line with current virological nomenclature practices, we distinguish between a formal taxonomic designation and a descriptive virus name. Accordingly, we refer to the genome sequences, viral lineage, physical isolates, and samples as *Peromyscus* coronavirus (PCoV) as previously described.

### Selection Pressure Analysis

Signals of site-specific positive, diversifying selective pressures were detected at 12 sites in the spike protein-coding sequence of *B. gravedinis* (Table 1). Notably, a cluster of sites subjected to positive selection were detected in the putative receptor binding domain (506 [p-value=0.03], 508 [p-value=0.01], and 517 [p-value=0.02]). Site 506 encodes for leucine, site 508 for phenylalanine, and 517 for serine. In a site-specific comparison of natural selection differences between the spike protein-coding sequence of *B. gravedinis* and its closest sister taxa, porcine hemagglutinating encephalomyelitis virus (PHEV), no sites were detected to be under statistically different natural selection regimes (Figure 2).

**Figure 2:**
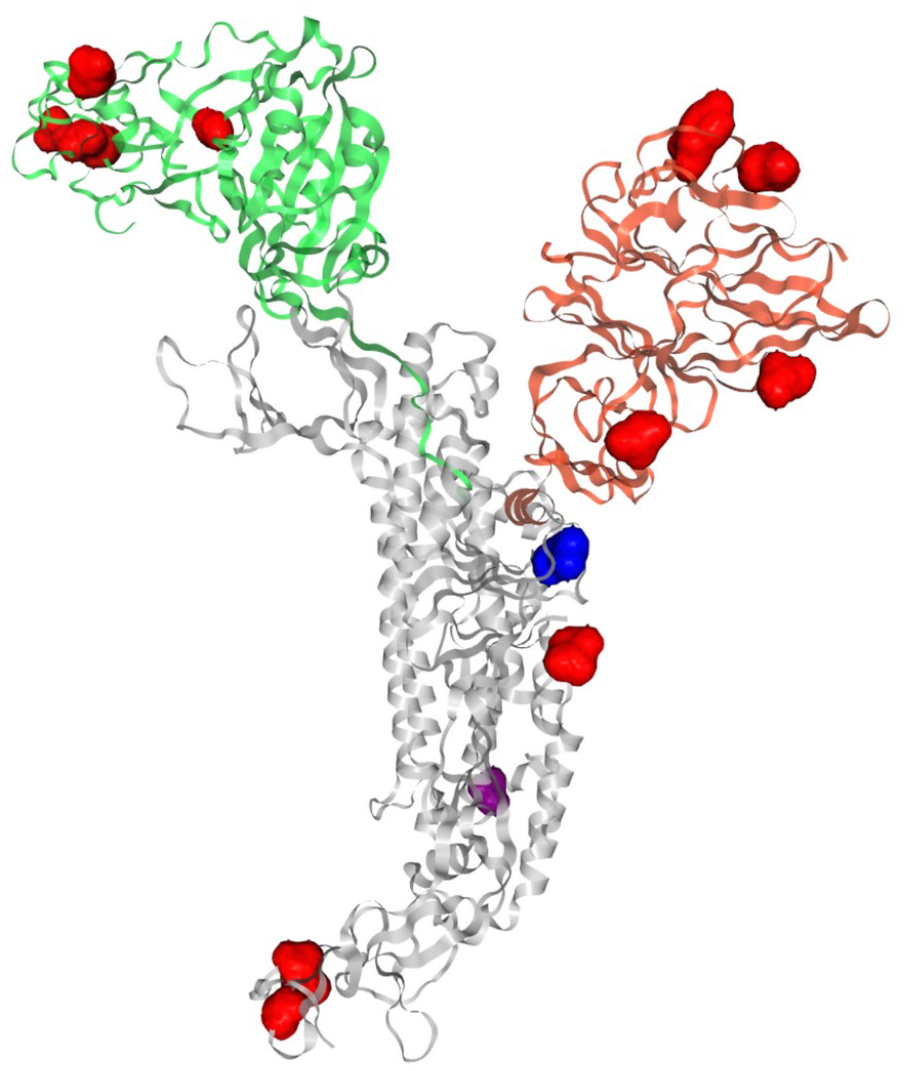
PHEV PDB 9H3J monomer displayed with domains NTD (red), S1 RBD (green), S1/S2 furin cleavage domain (blue), and S2 domain colored (yellow), with positively selected sites in *B. gravadinis* mapped in red.

### Geographic Distribution of Samples

Saliva swabs and fecal samples were tested for the presence of CoV RNA using a previously described PanCoV PCR assay. From 66 individuals collected in 2023 in Queens and Staten Island, CoV RNA was detected in the feces of 3 white-footed mice collected in Cunningham Park, Queens. No CoV RNA was detected in samples collected in Staten Island. Of 67 fecal samples collected in 2024 in Queens, CoV RNA was detected in one sample from a mouse in Alley Pond, Queens. No CoV RNA was detected in saliva swabs.

### Phylogeny of North American Rodent Species

A phylogeny was generated of North American rodents using the mitochondrial cytochrome b (cytb) gene to assess the evolutionary histroy and phylogenetic closeness of these species, especially between *P. leucopus* and *P. sonoriensis* (previously reported to be a potential host of *B. gravadinis*^9^). This phylogeny of North American rodents includes the most comprehensive number of species of this group to date. There was good support for the current families and their relationships to one another with the exception of the kangaroo rat family *Heteromyidae* (Figure 3). As previously reported, *Heteromyidae* appears paraphyletic without the inclusion of the pocket gopher family Geomyidae^20^. Similarly, most genera were well-supported with the notable exception of *Peromyscus* (deer mice) itself which was paraphyletic with respects to other new world mice genera in the subfamily *Neotominae* (*Cricetidae*; deer mice family), namely *Reithrodontomys, Megadontomys, Osgoodomys*, and *Habromys* (Figure 3). These results support previous findings suggesting a revision of Neotominae is necessary^21^. This analysis shows that *P. gossypinus, P. purepechus, P. gambelli*, and *P. maniculatus* form a clade with both known *B. gravadinis* host species: *P. leucopus* and *P. sonoriensis* (Figure 3).

**Figure 3:**
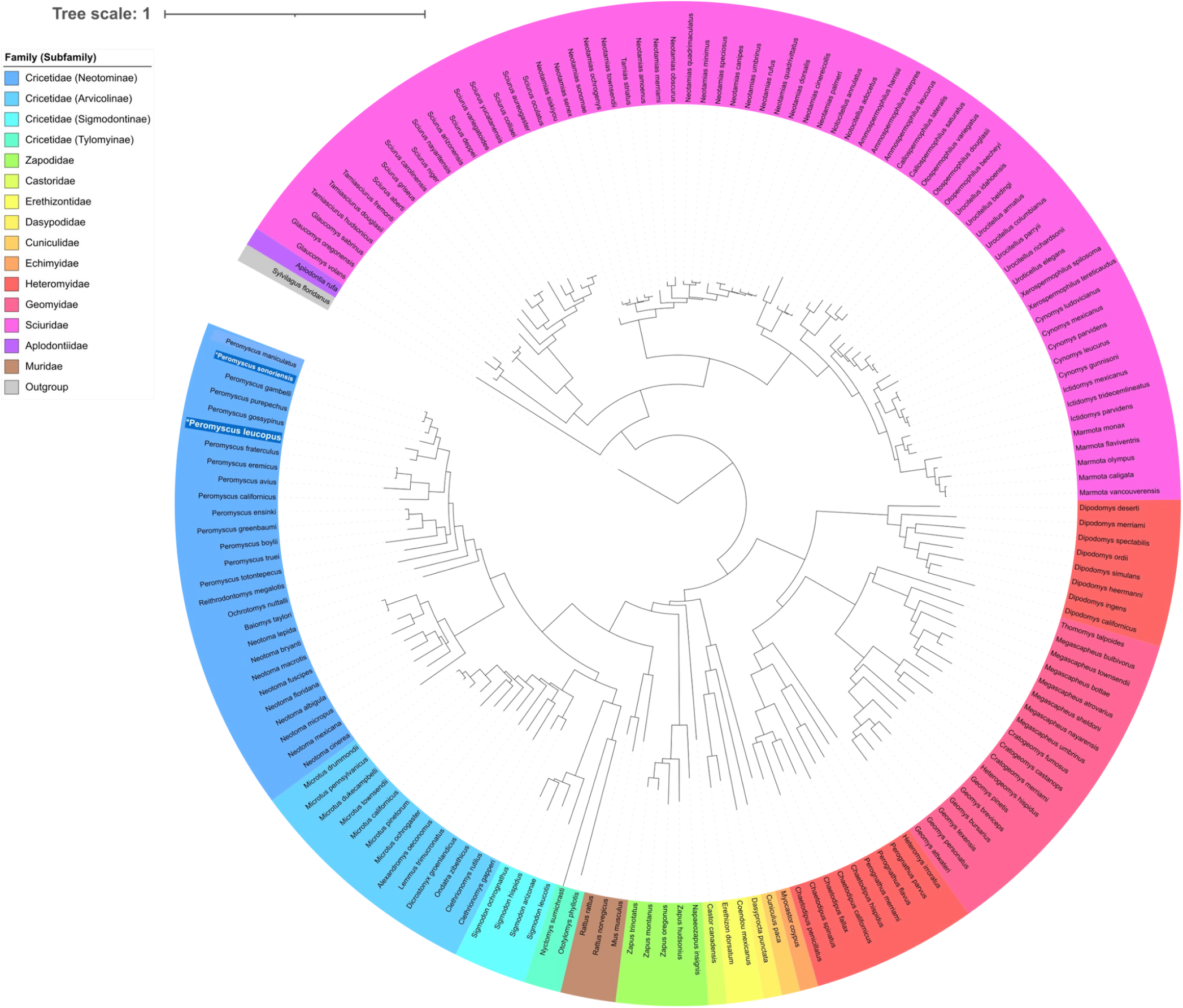
Maximum-likelihood phylogeny of North American rodents that appear in species distribution modeling. Known *B. gravadinis* hosts *P. leucopus* and *P. sonoriensis* are indicated with an asterisk and white font. Tip name color indicates species family, or in the case of the *Peromyscus* family *Cricetidae*, subfamily.

### Range shift of P. leucopus

Using species distribution models informed with 19 CHELSEA layers, the probable future range of *P. leucopus* is predicted to shift slightly northward under a middle-of the road climate scenario for the mid-21^st^ century (SSP 2-4.5) (Figure 4). The relative feature importance for the SDMs indicate that Mean Temperature of Warmest Quarter (BIOS 10) and Precipitation Seasonality (Coefficient of Variation) were both highly important in in prediction (Supplemental Figures 1 & 2).

**Figure 4:**
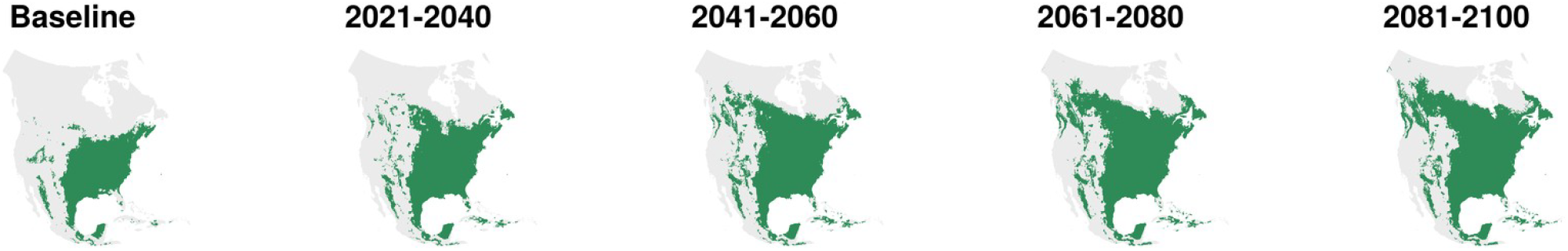
Future range shifts of *Peromyscus leucopus* under a middle-of-the-road global warming projection.

### Weighted Species Distribution Models based on Host Phylogenetic Distance

To predict where similar *B. gravedinis* genomes may be found in *P. leucopus* or other species, weighted species distribution models were generated and the biodiversity dose and uncertainty were calculated (Figure 5). The results of weighted species distribution modeling, corresponding to different possible transmission regimes among rodent species in North America, suggests that there are three hotspots where circulation is likely across all transmission regimes. First, a region starting with the Delaware river watershed extending northwards along the coast to Nova Scotia and extending inland into the Great Lakes Watershed was present under all regimes tested. Variation in this hotspot was observed particularly between the linear regime versus the exponential or gaussian regime, with the linear regime predicting a more consistently high dose in the Great Lakes watershed and further inland to the south, whereas this region appeared more fragmented in both the gaussian and exponential regimes. The second is the western coast of North America starting from the Salish sea extending down to the start of the Baja California peninsula. Across all transmission regimes, this region did not vary significantly between different transmission regimes. The third region is the southern coast, starting after the gulf of California (about where Baja California ends on the mainland side) extending south to the southern pacific tip of Oaxaca. This region appeared to extend further inland in the exponential and gaussian regimes, whereas the corresponding region was more coastally restricted in the linear model.

**Figure 5:**
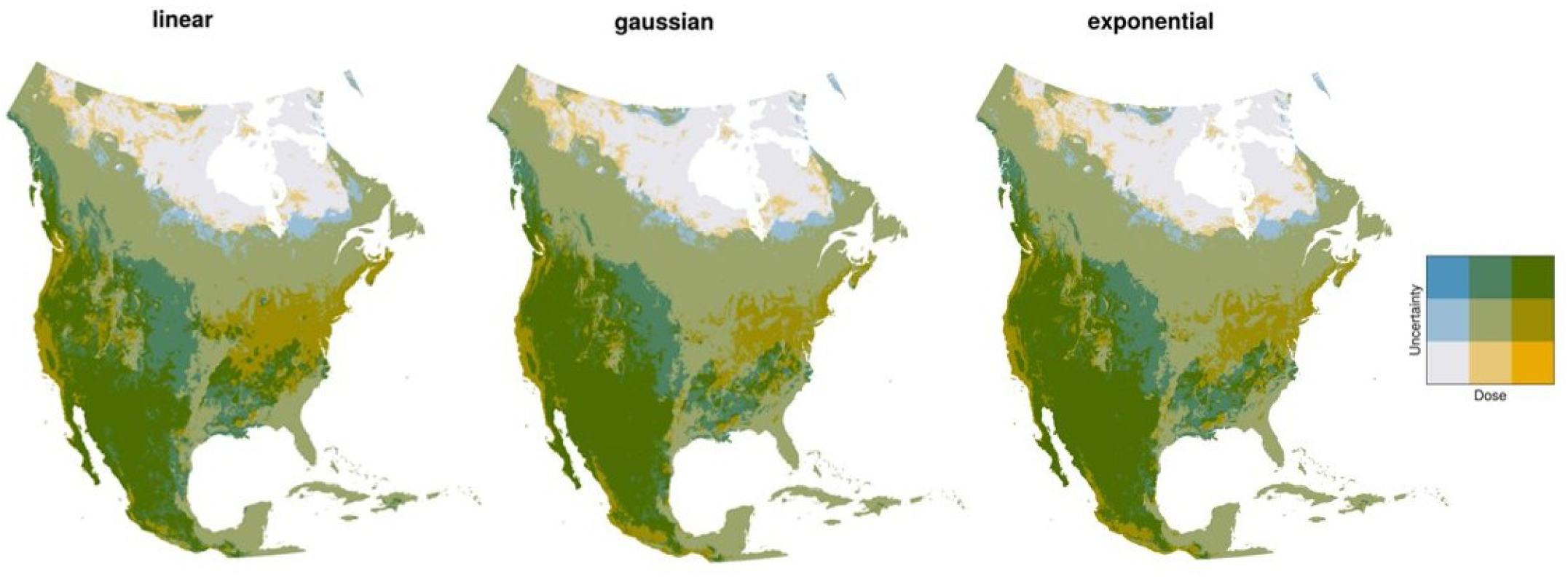
“Dose” of *Peromyscus*-origin *B. gravadensis* in North America under different possible transmission regimes

### Changes in Distribution due to Climate Change

Under a middle of the road climate change scenario SSP2-4.5, all three models converge on a picture of a new hotspot appearing south of the Hudson Bay with a major hotspot in the Cree lands on the eastern coast of Baie James (Figure 6). In the linear model, the western coast of Newfoundland, Gaspésie and Île d’Anticosti are also predicted to experience substantial gains in suitability for *B. gravadinis*. In all three models, a region south of the Great Lakes extending northwestwards into the plains is predicted to experience the greatest loss of suitability. In both the exponential and gaussian models, the northeastern Atlantic coast including the New York City Metropolitan area and Connecticut are predicted to experience loss whereas this is not present in the linear model.

**Figure 6:**
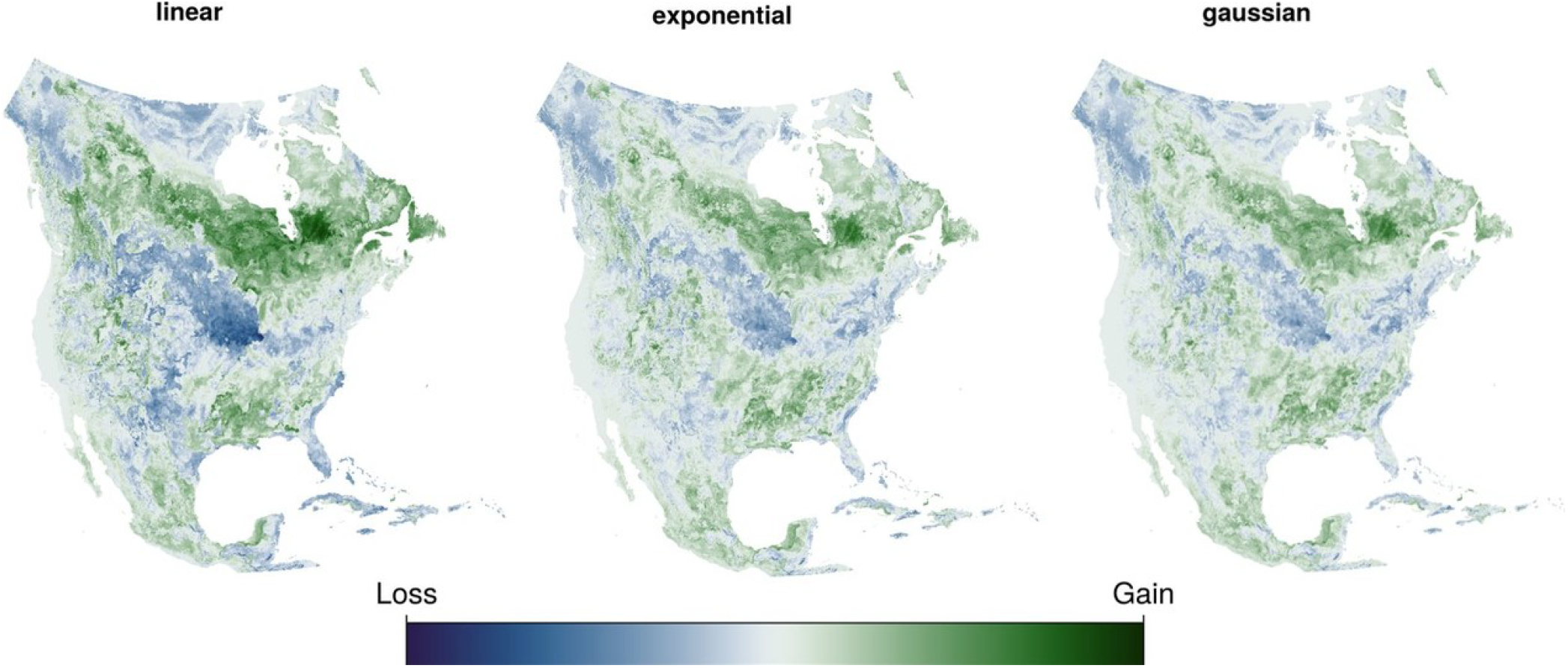
Climate Induced Geographic Shifts in Weighted Species Distribution Models

## DISCUSSION

Rodents (order *Rodentia*) comprise over 2,200 species and represent the most speciose mammalian order^22^. Their wide geographic distribution, adaptability to urban environments, and potential for high-density populations make them prime candidates for supporting spillovers to humans^2,23–27^. Several major viral pathogens with significant human health impacts have emerged from rodent hosts, including hantaviruses, arenaviruses, and most recently, mpox virus, which has been linked to African rodent species^28–31^.

Within the genus *Betacoronavirus*, the subgenus *Embecovirus* contains several human coronavirus members (e.g., OC43, HKU1) that are thought to have originated from rodents or ungulates^32,33^. Rodent-associated *Betacoronavirus* species have been previously detected in North America, but their diversity, evolutionary history, and zoonotic potential have not been addressed in great depth^9,11,13,34,35^. However, recent work has shown that rodent reservoirs can harbor non-rodent origin coronaviruses such as SARS-CoV-2^13,35,36^.

The phylogenetic position of the genomes sequenced in this study fall within the *B. gravedinis* species. There is a high level of similarity between genomes across multiple sites and over time and the amplification products were obtained only from rectal swabs which was consistent with previous reports of the virus only being present in colon or rectal tissue, except for equivocal reports in lung tissue in the case of 2 animals^9,13^. Positive samples were obtained during the summer which is consistent with the 2022 study reporting detection of *B. gravadinis* in *P. leucopus* in Connecticut, however this could equally be a result of sampling bias of trapping seasons occurring in summer months. Future studies may investigate if there are seasonal or climactic factors that influence viral circulation such as seasonal pulses following precipitation. Additionally, future field studies may consider attempting viral isolation on feces versus saliva, or other infected tissues to confirm which materials the virus can persist in.

The 3 sites subject to positive selection in the RBD of spike consisted of a leucine, phenylalanine, and serine (sites 506, 508, and 517, respectively). Selection at these sites could be indicative of receptor affinity, structure stabilization, immune escape, or implicated in another feature. Our selection analysis identified several regions under positive pressure, including in spike, suggesting ongoing adaptation within the rodent host. Notably, there were no statistical differences detected in selection regimes between rodent-origin *B. gravedinis* genomes and the sister lineage PHEV. The absence of lineage-specific selection could suggest that these proteins traverse a permissive evolutionary landscape, where common structural foundations facilitate the exploration of diverse phenotypic space^37^. This confirms what has been previously reported in the literature where OC43-like viruses within *B. gravidinis* have been described as “promiscuous” coronaviruses, capable of infecting a diverse set of mammalian hosts including humans, deer-mice, deer, dogs, giraffes, antelope, camels, horses, and cows^9^. Future wet-lab studies will be crucial to elucidate the impacts of mutations at these positions, and to experimentally determine the tissue tropism or receptor.

Rodents in the genus *Peromyscus* are established reservoirs of well-characterized zoonoses with mapped geographic ranges. *Peromyscus leucopus* is the primary reservoir of *Borrelia burgdorferi* across the northeastern United States and the Great Lakes region^38^ and a known host of *Orthohantavirus sinnombreense* (Sin Nombre hantavirus, strain New York-1)^39^, and *P. sonoriensis* is the principal reservoir of Sin Nombre hantavirus across western North America^40^. In 2020, a genome called “BetaCoV/Peromyscus maniculatus/Utah/238640/2020” was reported in a related *Peromyscus* species in Utah clustering in the same lineage of the *Embecovirus* genus^9^. It was reported that 3 additional animals were positive based on an observed amplification product of similar size seen on the gel. In summer 2022, *B. gravadinis* genomes PCoV1 and PCoV2 were detected in *P. leucopus* at two sites in Connecticut^13^.

The results of the weighted SDMs generated in this study recover the same regions as the previous detections of *B. gravadinis* in *Peromyscus* rodents. Interestingly, the output of the model reproduces the known geography of other pathogens (including *B. burgdorferi* and *O. sinnombreense* strain NY-1) that share the same reservoir. The most uncertainty between the predictions can be accounted for by how contiguous or fragmented the predicted hotspot is. Future field studies may consider focusing on these three current predicted regions to obtain more sequences or to isolate the virus.

As the climate warms, the risk of future spillover events is thought to increase^41^. When similar predictions are made for a future North America, accounting for warming climate, there is an additional hotspot that appears in Jamésie Cree land east of James Bay in all three transmission regimes tested and extends broadly into all Cree Lands bordering Hudson Bay. This hotspot in Cree Lands can be taken into account in order to safeguard the health of people and animals living in those lands if a risk to human health is established for this virus.

## METHODS

### Animal Trapping and Sample Collection

*Peromyscus leucopus* (white-footed mice) were trapped using Sherman live traps and sampled from the beginning of August - early September 2023 and 2024 at parks across New York City (Table 2). Traps were baited with oats and peanut butter, and cotton was provided as nesting material. Traps were checked the following morning from 6:00 am - 12:00 pm and all trapped animals were sampled and released back to the trapping location without the use of sedation. Samples included collection of feces and oral swabs from each individual. Feces and oral swabs were stored separately in 1.5-mL screw top microcentrifuge tubes with 500ul of DNA/RNA Shield (Zymo Research). Samples were transported on ice and stored at 4°C prior to extraction. Capture and sampling of *P. leucopus* was performed under New York State Department of Environmental Conservation permit #2704 and Columbia University IACUC protocol #AC-AABM9557.

### Pan coronavirus screening and sequencing

RNA was extracted from feces and saliva swabs as previously published^42^. The obtained RNA was reverse transcribed using the LunaScript RT SuperMix (New England Biolabs, NEB). The cDNA from feces and saliva swabs was screened with a pancoronavirus assay using previously published primers targeting a conserved region of the RNA-dependent RNA polymerase (RdRp)^43^. For the samples collected in 2023, all the feces and saliva swabs were tested. For the samples collected in 2024, only those that tested positive in feces were screened for CoV in the saliva samples. The PCR process was carried out as previously published^42^. Positive samples were cleaned using the QIAquick PCR Purification Kit (QIAGEN), and the final DNA concentration was measured using the Qubit 4 Fluorometer (Thermo Fisher Scientific) using the dsDNA Broad Range Quantification Kit (Thermo Fisher Scientific). PCR library preparation and sequencing using the MinION (Oxford Nanopore Technologies) were done as previously described^44^.

### Genome Assembly and Phylogenetic Analysis

The whole genome of confirmed CoV-positive samples was sequenced using a primer scheme (Supplementary Table 1) based on the sequences of closely related CoVs found after BLAST of the partial RdRp sequences obtained during the screening process (accession numbers OR735432 – 34, OR613113). The primer scheme was designed using PrimalScheme^45^. PCRs were carried out using the Phusion Hot Start II High-fidelity PCR Master Mix (Thermo Fisher Scientific), using an annealing temperature of 60 °C. PCR products were cleaned using 1.8X of AMPure XP Beads (Beckman Coulter Life Sciences). Each PCR pool was sequenced in the MinION as previously described^44^. The super-accurate model was used for basecalling to obtain reads with a minimum Q score of 20. During the basecalling process, the barcodes were trimmed on both sides of each sequence. Primer sequences were trimmed from the high-quality sequences obtained using the Trim Ends option in Geneious Prime 2023.2.1 (Biomatters). Trimmed sequences were mapped against the most closely related sequences (accession numbers OR735432 – 34, OR613113) using the Geneious mapper with custom sensitivity and three iterations in Geneious Prime 2023.2.1 (Biomatters). A consensus was generated using a threshold of 75%. The consensus sequences obtained were uploaded to GenBank under accession numbers PZ468949 to PZ468950.

To investigate the evolutionary relationships of the newly identified virus with other known rodent coronaviruses, phylogenetic analysis was performed based on the RNA-dependent RNA polymerase (RdRp) region encoded by the NSP12 gene. Viral genomes for the two positive samples (SM009RE and SM004RE) were aligned against related betacoronaviruses (Table 1) and annotated using the NSP12 coordinates from Mouse Hepatitis Virus A59 (NC_048217). Nucleotide sequences corresponding to the NSP12 region were extracted from each genome in Geneious (Dotmatics) and aligned using Clustal Omega.

Whole-genome nucleotide sequences of 24 coronaviruses were compiled and aligned to reconstruct their evolutionary relationships. Each genome was renamed using its GenBank accession number to ensure consistent downstream handling. The sequences were aligned using MAFFT v7.505 with the --auto option for automatic strategy selection and default gap penalties. The resulting multiple sequence alignment contained 34,596 positions. Phylogenetic inference was performed using IQ-TREE v2.2.6 under the GTR+G substitution model, which was selected automatically based on model testing.

Ultrafast bootstrap approximation with 1000 replicates (-B 1000) was conducted to assess node support. The resulting tree was rooted using the outgroup method and exported in Newick format as a .treefile.

### Phylogenetic Relatedness of North American Rodents

In order to assess the phylogenetic closeness of North American rodents to known *B. gravadinis* hosts *Peromyscus leucopus* and *P. maniculatus*, a phylogeny and distance matrix of North American rodents was constructed. North American rodents were defined as any species with a portion of their range in Canada, the United States, or Mexico. Species ranges and taxonomy were determined using The Mammal Diversity Database by the American Society of Mammologists resulting in 431 target species. Mitochondrial cytochrome b (cytb) gene sequences for each target species were selected and downloaded from NCBI GenBank based on sequence length and quality. Of the 431 target species, 49 had no cytb sequences available, leaving a complete dataset of 384 species including the outgroup, *Sylvilagus floridianus*. Sequences were also aligned in MAFFT v.7.505 using the --auto option and default gap penalties. The phylogeny was inferred using a GTR+G maximum-likelihood substitution model with non-parametric bootstrapping as implemented in raxml-ng v.2.0.1 using the --all option.

The phylogenetic distance between each species pair was assessed using both an outgroup rooted and unrooted tree topologies to form the distance matrices used in downstream analyses. This matrix was created using the cophenetic.phylo() function in the R package ape.

### Weighted Species Distribution Modelling

To predict what other rodent species in North America might be susceptible or harbor *B. gravedinis*, phylogenetic distance was used as a surrogate measure for *“*hostness” based on the assumption that probable hosts would be more frequent among closely related rodent species. Given the low prevalence of finding these genomes in white-footed mice, we developed models for three different possibilities.

First, that white-footed mice or a very closely related species is the primary host and shares the virus with close phylogenetic relatives. Secondly, that the virus is a host generalist and capable of infecting many different hosts but with a preference for infecting white-footed mouse or a closely related species. Thirdly, that the virus is capable of infecting a broad range of rodent species and that any one of the species it infects could be considered a primary host.

Weights are computed as follows: for a focal species i, the weight for species j is computed as its relative “closeness” to species i. Species are first assigned a score s_j based on one of several distance functions. The score using linear distance function is computed as sj = M - d{ij}, where M is the maximum distance between i species in the tree. For the exponential distance function, sj = exp(- \alpha * d{ij}). And for the Gaussian distance function, sj = exp(-d{ij}^2/(2\sigma^2). Here, both \alpha and \sigma = 1. Then, weights are computed from these scores as wj = d{ij} / sumk d{ki} for all species j.

### Natural selection analysis of Spike protein

Codon-aware alignments were generated from spike protein coding sequences; 6 *Betacoronavirus I* (3 sequences reported herein, and 3 reported from another study) and 100 porcine hemagglutinating encephalomyelitis virus (PHEV) spike sequences, as PHEV is sister taxa to *Betacoronavirus I*. following the procedure available at the Github repository <u>Codon-MSA</u> (github.com/veg/hyphy-analyses/tree/master/codon-msa). Briefly, in-frame nucleotide sequences were translated, aligned with multiple alignment using fast Fourier transform v7.471, and then mapped back to corresponding nucleotide sequences^46^. A single copy of identical sequences was retained. To prepare the hypothesis testing with phylogenies software suite (HyPhy v2.5.93^47^), the genetic algorithm for recombination detection (GARD^48^)and partitioned into recombinant-free partitions (RFP) accordingly. A phylogeny was then inferred for each RFP with RAxML-NG v. 1.2.2^49^. The 6 *Betacoronavirus I* branches were labeled with phylotree.js^50^, the mixed effects model of evolution (MEME^51^) was used to detect signatures of site-specific patterns of positive, diversifying selection across the 6 *Betacoronavirus I* (3 sequences reported herein, and 3 reported from another study).

## Notes

### Competing Interest Statement

The authors have declared no competing interest.

