## Supplement for "Genomes of *Betacoronavirus gravedinis* from white-footed mice in New York City and a phylogenetically weighted model of its probable distribution in North America"

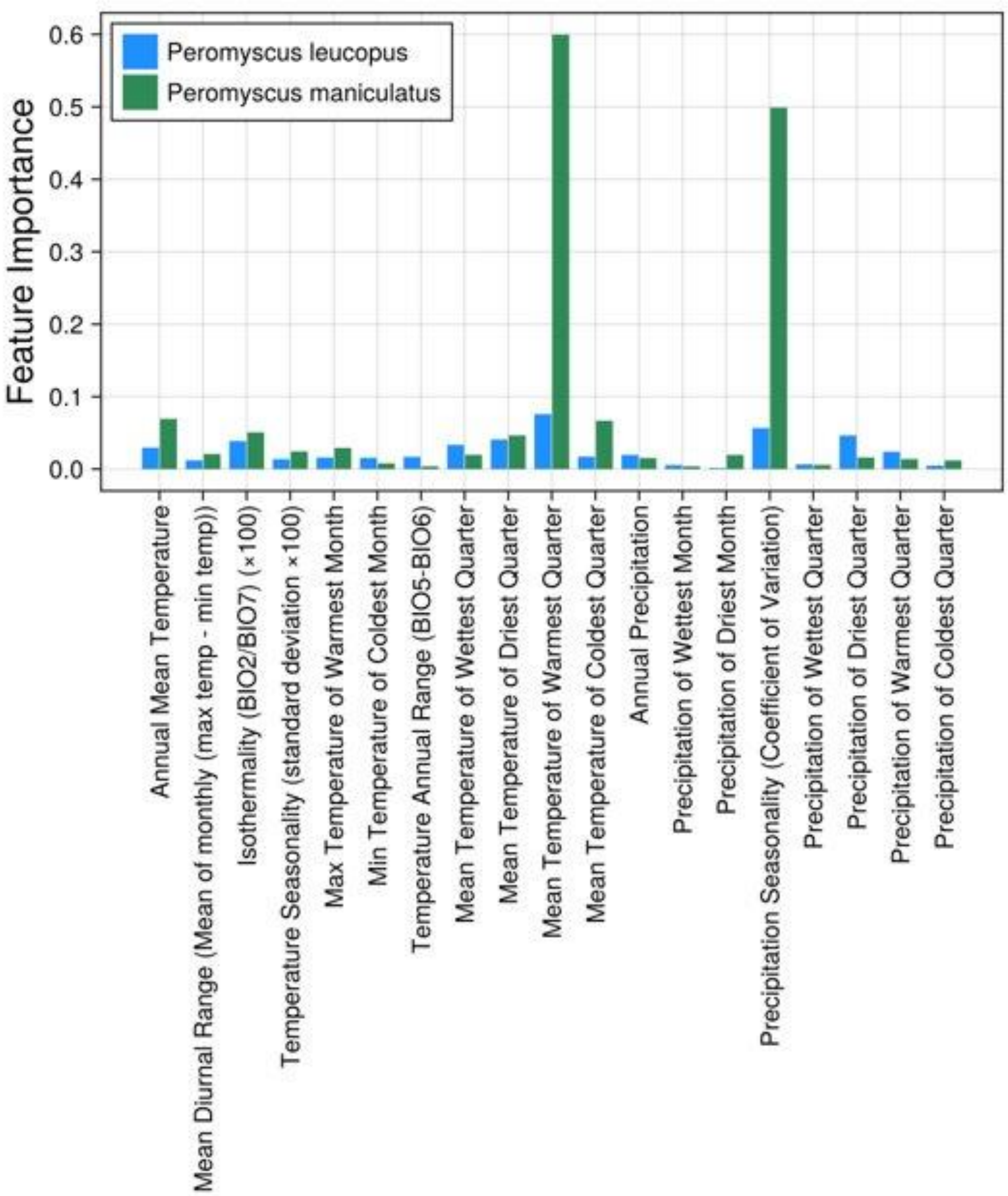

**Supplemental Figure 1:** Relative feature importances for *Peromyscus leucopus* and *maniculatus* in SDMs.

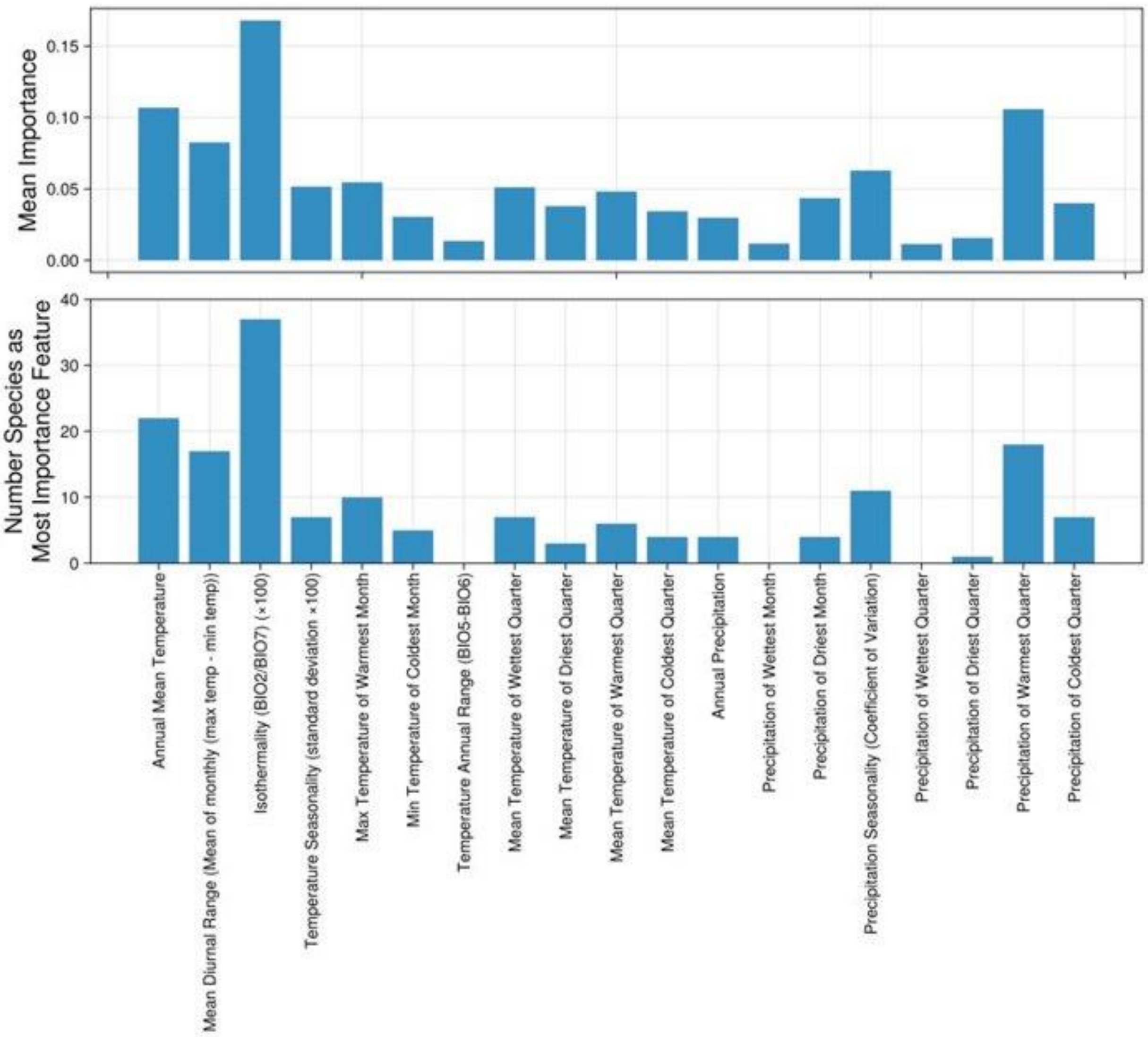

**Supplemental Figure 2**: Relative Feature importances for all rodent species used in SDMs.

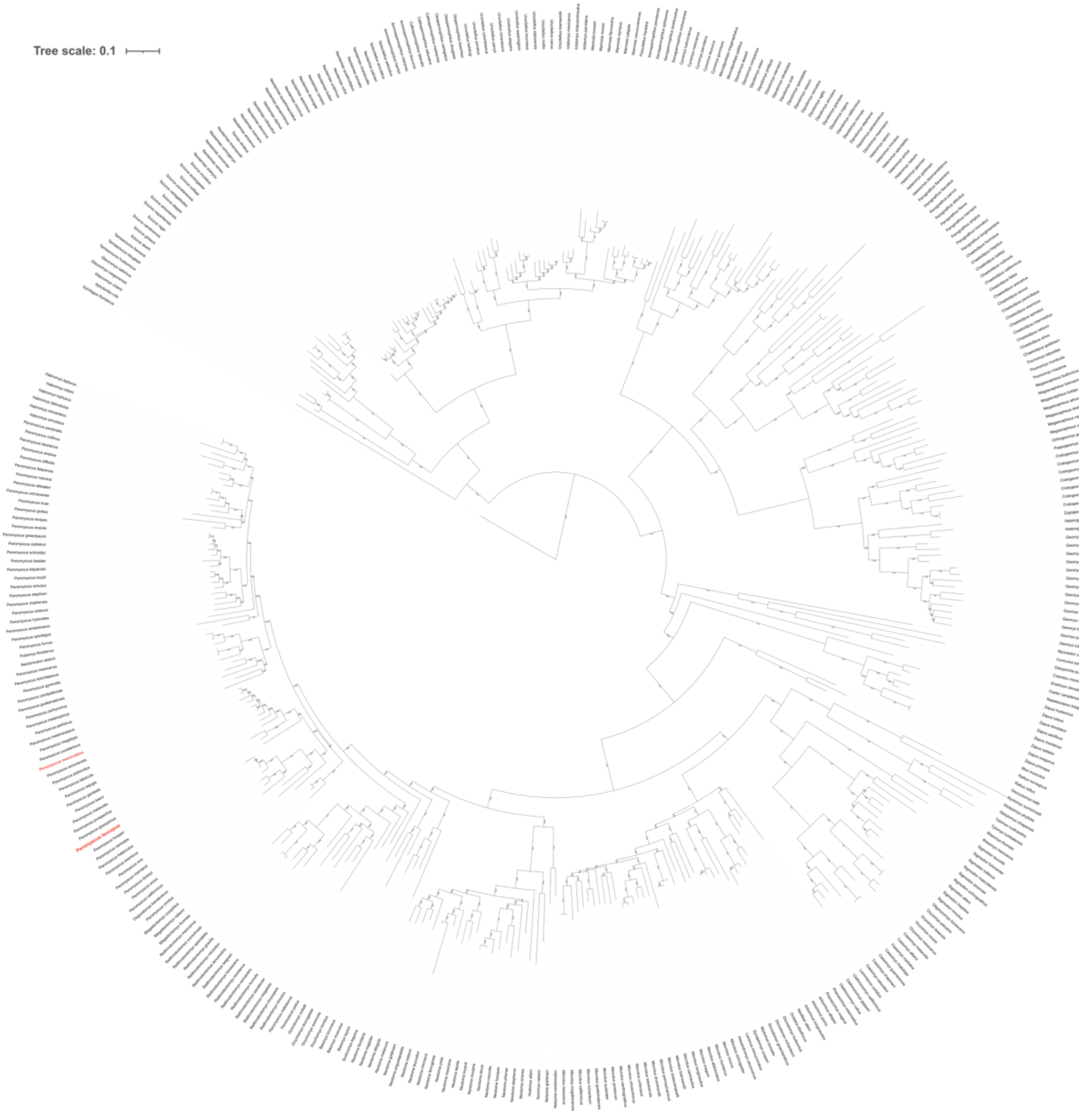

**Supplemental Figure 3.** Phylogeny with bootstrap values of all North American rodents with available CYTB sequences.

**Supplementary Table 1.** Primers used for whole-genome sequencing of the coronavirus detected in the samples.

| Primer ID | Primer sequence (5' - 3') | Primer pool |
|---|---|---|
| BCoV_1_F | TCCCTGTAGTCTATGCTTGCGA | 1 |
| BCoV_1_R | CCATCCAACAGGAATGGCATAAAAC | 1 |
| BCoV_2_F | CTTGGTTTGTCAGATGTGTGGC | 2 |
| BCoV_2_R | TCGGCAACATAACCACCTTGTG | 2 |
| BCoV_3_F | CGTTCAGTTTTGGCAGTGATGC | 1 |
| BCoV_3_R | ACAACAACCTGCCACTTAGAACA | 1 |
| BCoV_4_F | GGTTTGTGCCTGTTGGTGAAAG | 2 |
| BCoV_4_R | CCACGAGCAACTACACTCCAAT | 2 |
| BCoV_5_F | AGCCTGTTGTTAATGTGGTTAAAGTG | 1 |
| BCoV_5_R | GGTGTGGTTAAAACAGAGACATTAGC | 1 |
| BCoV_6_F | CTAAGCATGTGCAGGGTAACGT | 2 |
| BCoV_6_R | GCAGAACATATCACATGTCTGGGA | 2 |
| BCoV_7_F | TGCTTTACCAACCACCTACAGC | 1 |
| BCoV_7_R | GGCACACCTCCAATACCCAAAA | 1 |

| | | |
|---|---|---|
| BCoV_8_F | GCTGTTATTGGGGTTTGTTTTCCC | 2 |
| BCoV_8_R | CATCCCGCCAAAATCCACAAAC | 2 |
| BCoV_9_F | GTCTGTATTTATTGCCGCGCAC | 1 |
| BCoV_9_R | ATGCCTCATAGTAAAGCCTGGC | 1 |
| BCoV_10_F | AAGGGCCTGCTTAAAGAGGGTA | 2 |
| BCoV_10_R | GTACGGAGATCCAGTACAAGATTGT | 2 |
| BCoV_11_F | CCTCTTCTTTGCTGCAAGTGTTG | 1 |
| BCoV_11_R | CCAATACCAAGACATACGGCCA | 1 |
| BCoV_12_F | CGCTTCAATGTTGCTATTACTCGAG | 2 |
| BCoV_12_R | TCCTCCTTATCGATCTTAGCCACA | 2 |
| BCoV_13_F | TGCTGAAGAGTATCGTGAGTACCT | 1 |
| BCoV_13_R | CCGTCCAAAATGCAAAATACCCC | 1 |
| BCoV_14_F | TGGAAGCCTGGTTATTCTATGCC | 2 |
| BCoV_14_R | CAATCTGAACGACTGTCACCGA | 2 |
| BCoV_14.2_R | CAATCTGAACGACTGTCACCRA | 1 |
| BCoV_15_F | CCCACATCCTTGGAAGATGCAT | 1 |

| | | |
|---|---|---|
| BCoV_15_R | ACGCATCATGCAACCTAGTACC | 2 |
| BCoV_16_F | TTGGGTCACACCTCTCACTAGT | 2 |
| BCoV_16.2_F | TTGGGTCACACCTCTCACTART | 1 |
| BCoV_16_R | AGCTACTTGCAACTGTGTAGTGTC | 1 |
| BCoV_17_F | GAGTTTATTCAGACGAGCTCTCCT | 2 |
| BCoV_17_R | CAGAGGTAAAACTCACCACGCT | 2 |
| BCoV_18_F | AGAAATGTGGTGGTTGTTGTGATG | 1 |
| BCoV_18_R | ACCTTGAATGTAGAGGTGGCCA | 1 |
| BCoV_19_F | GTGGCCATTATTATGTGGATTGTGT | 2 |
| BCoV_19_R | AGAGAGTGCCTTATCCCGACTT | 2 |
| MBCoV_F* | TGTTGACGTCACCAAACACA | -- |
| MBCoV_R* | ATAATGCAAGCACCACGAGC | -- |
| MBCoV2_F* | ATGTTAGCAGTGAGTGGCCT | -- |
| MBCoV2_R* | ACAACCAAAAGAGTCTGCCTG | -- |
