## Supplementary material for "Genomes of *Betacoronavirus gravedinis* from white-footed mice in New York City and a phylogenetically weighted model of its probable distribution in North America": Tables

| Codon Position Mapped to sequence sm009WFM | P-value |
|---|---|
| 25 - TS**S**IN | 0.03 |
| 40 - TE**V**VD | 0.01 |
| 186 - WH**W**DT | 0.04 |
| 204 - DV**N**AD | 0.04 |
| 506 - VR**K**CF | 0.03 |
| 508 - KC**F**DY | 0.01 |
| 517 - EC**S**CW | 0.02 |
| 564 - PC**T**CT | 0.04 |
| 572 - FV**S**WG | 0.02 |
| 747 - DY**V**TA | 0.04 |
| 1167 - GD**R**GI | 0.01 |
| 1203 - VV**V**MS | 0.02 |

**Table 1:** Positively selected sites in *B. gravedinis* as identified by MEME. For each site, the selected amino acid is bolded and flanked by two upstream and two downstream residues to provide structural and sequence context.

| Borough | Park | Number of individuals sampled | |
|---|---|---|---|
| | | 2023 | 2024 |
| Queens | Alley Pond | 4 | 6 |
| | Cunningham | 17 | 29 |
| | Forest Park | 11 | 18 |
| | Kissena Park | 8 | 14 |
| Staten Island | Blue Heron | 1 | -- |
| | Conference House | 9 | -- |
| | Lemon Creek | 10 | -- |
| | Long Pond | 1 | -- |
| | Mount Loretto | 5 | -- |
| | Total | 66 | 67 |

**Table 2**. Sample collection of white-footed mice in Queens and Staten Island. For each individual, feces and saliva swabs were collected.
